# A Meta-Analytic Single-Cell Atlas of Mouse Bone Marrow Hematopoietic Development

**DOI:** 10.1101/2021.08.12.456098

**Authors:** Benjamin D. Harris, John Lee, Jesse Gillis

## Abstract

The clinical importance of the hematopoietic system makes it one of the most heavily studied lineages in all of biology. A clear understanding of the cell types and functional programs during hematopoietic development is central to research in aging, cancer, and infectious diseases. Known cell types are traditionally identified by the expression of proteins on the surface of the cells. Stem and progenitor cells defined based on these markers are assigned functions based on their lineage potential. The rapid growth of single cell RNA sequencing technologies (scRNAseq) provides a new modality for evaluating the cellular and functional landscape of hematopoietic stem and progenitor cells. The popularity of this technology among hematopoiesis researchers enables us to conduct a robust meta-analysis of mouse bone marrow scRNAseq data. Using over 300,000 cells across 12 datasets, we evaluate the classification and function of cell types based on discrete clustering, *in silico* FACS sorting, and a continuous trajectory. We identify replicable signatures that define cell types based on genes and known cellular functions. Additionally, we evaluate the conservation of signatures associated with erythroid and monocyte lineage development across species using co-expression networks. The co-expression networks predict the effectiveness of the signature at identifying erythroid and monocyte cells in zebrafish and human scRNAseq data. Together, this analysis provides a robust reference, particularly marker genes and functional annotations, for future experiments in hematopoietic development.

**Key Points:**

- Meta-analysis of 9 mouse bone marrow scRNAseq identifies markers for cell types and hematopoietic development
- Characterize a replicable functional landscape of cell types by exploiting co-expression

## Introduction

The hematopoietic lineage is one of the most highly studied lineages in all of developmental biology. Classically, cell types are identified by Fluorescent Activated Cell Sorting (FACS). For example, the Long Term Hematopoietic Stem Cell (LT-HSC) is identified by CD34low, Flt3-, and TpoR+ expression. The role of a progenitor is to produce differentiated cells, and the function of a specified progenitor cell type is defined by the potential to differentiate into a specific lineage ^1^. The Multipotent Progenitor 4 (MPP4) is heavily biased toward differentiation into the lymphoid lineage ^2^. Importantly, the discovery and functional annotation of cell types are dependent on the modality of data. FACS and lineage potential are not the only such methods ^3–5^. The advent of single-cell RNA sequencing (scRNAseq) allows for the classification of the hematopoietic lineage from an entirely new data modality. Using gene expression to characterize cell types gives us a new opportunity to identify the gene regulatory programs important to hematopoietic lineages.

A clear understanding of hematopoietic development is central to aging and cancer research. The bias towards the development of myeloid cells instead of the lymphoid lineage is a major molecular signature of aging ^6–8^. Additionally, some hematological cancers can be viewed as misregulation or stalled development of myeloid cells, leading to a class of therapeutics known as differentiation therapy ^9,10^. Identifying the changes in gene regulation that cause lineage bias or developmental stalling is crucial to perturbing these systems back into a healthy state. An atlas that describes cell types involved in healthy hematopoiesis, and characterizes the function for each cell type using scRNAseq will serve as a critical reference for translational research.

The rapid development of scRNAseq technology creates the opportunity to build a robust atlas of hematopoietic cells in the bone marrow. Multiple studies publish individual atlases of hematopoietic development, but they do not integrate information from other published datasets ^11,12^. Replicability across many datasets resolves some technical limitations of individual scRNAseq datasets, creating a more robust atlas ^13,14^. The most comprehensive and robust cell atlases rely on meta-analysis across many scRNAseq datasets to characterize replicable cell types ^15–18^. After identifying the present cell types, characterizing their functions can be done by evaluating the signatures that define the cell types for known processes and pathways ^19–21^.

In this work, we build a comprehensive mouse hematopoietic cell atlas by integrating and labeling over 300,000 cells from 14 datasets. We identify robust gene regulatory signatures using multiple perspectives of the data. Two bone marrow datasets from the Tabula Muris consortium and the semi-supervised machine learning algorithm scNym are used to label and integrate 12 datasets of mouse bone marrow data ^22,23^. We identify robust markers for each cell type and learn functional annotations using the Gene Ontology. Labeling cells based on genes that traditionally serve as cell surface markers identifies a latent lineage potential signature. Pseudotime analysis finds signatures associated with the development of the monocyte and erythroid lineages. Co-expression and scRNAseq from zebrafish and human samples evaluates the conservation of lineage-associated signatures in 21 species. We present a replicable view of hematopoietic development in the mouse bone marrow, that complements the FACS and lineage potential-based perspective of hematopoietic development.

## Methods

### Data preprocessing

Data were downloaded for each dataset based on the info provided by their publication. For a detailed explanation, see the code for each dataset in the Github repository.

### Integration using scNym

Data was normalized to logTPM (normalize_total=1e6) as per the requirements for scNym. A column in the Anndata object was created that had the cell type labels from the two tabula muris datasets and the placeholder “Unlabeled” for cells from all other datasets. Additionally, we included a column in the obs data that denoted the batch. When training and testing the model we use batch as the domain. The output layer, consisting of 256 features was used as the input to UMAP. All of this was run on a server with a Nividia Tesla V100 GPU and the UMAP was done using the Nvidia rapids library.

### Marker identification and enrichment

We use the MetaMarkers package in R to compute cell type markers for both the scNym labeled cell types (Figure 2) and in silico sorted cell states (Figure 3) ^24^. MetaMarkers computes differential expression using the Mann-Whitney test within each batch and then computes meta-analytic statistics to aggregate the statistics across batches. For the in silico analysis, we also use the score_cells, compute_marker_enrichment, and summarize_precision_recall functions to evaluate the identifiability and classification of cell states. Enrichment was done using the pyMN MetaNeighbor package and the mouse gene ontology.

### Pseudotime

Pseudotime was computed using monocle3 on each dataset ^25^. We tuned the parameters minimum_branch_length and rank.k to balance the complexity of the trajectory and the coverage of the lineages. We used the monocle2 differentialGeneTest function to calculate the genes associated with each lineage and with branching ^26^. For GO enrichment we used custom code for Fisher’s exact test (see GitHub) and the mouse gene ontology on the top 50 markers for each lineage.

### Cross-Species Co-expression

We evaluated the co-expression of orthologs to the lineage-associated gene lists for every species in CoCoCoNet with at least 5 orthologs for both lineages using EGAD ^27^. In the human data Pelin et al 2019 and zebrafish dataset Xia et al 2021 we scored the expression of the orthologs using the Scanpy score_gene_list functions.

### Data and code availability

The code for all analysis is available in the GitHub repository https://github.com/bharris12/hsc_paper and processed data is available on the FTP site ftp://gillisdata.cshl.edu/data/HSC_atlas/. The data can also be explored in the Shiny app at https://gillisweb.cshl.edu/HSC_atlas/

## Results

### Integration and Filtering of Datasets

We collected 12 published datasets that use high throughput scRNAseq methods to profile mouse hematopoietic progenitor cells ^11,28–33^. Not all of the original publications label every cell and each publication has unique rules for defining cell types. These two challenges make it unclear what cell types are shared across publications by looking at the published papers and associated metadata. It is preferable to have an integrated latent space with cells from all datasets for some analyses. After identifying the shared populations across the publications, we can evaluate the discrete and continuous models of cell types (Figure 1A). We use the tool scNym and the Tabula Muris bone marrow dataset, a high-quality reference dataset, to integrate and label the cell types from all of the studies. From 7 publications, we identified 12 sequencing batches that we refer to as datasets. Projecting individual datasets into a latent space using UMAP for Weinreb et al 2020 and Rodriguez-Fraticelli et al 2020 clearly shows technical variation between annotated batches, while Tikhonova et al 2019, despite annotating multiple batches, does not present strong batch effects (Supplementary Figure 1). We treat each of the batches in the 3 publications as individual datasets to avoid fitting to technical variation within some datasets. Projecting all the cells into a low dimension integrated UMAP space shows the clustering of cell types based on the scNym labels and consistent overlap for most of the datasets (Figure 1B-C).

**Figure 1:**
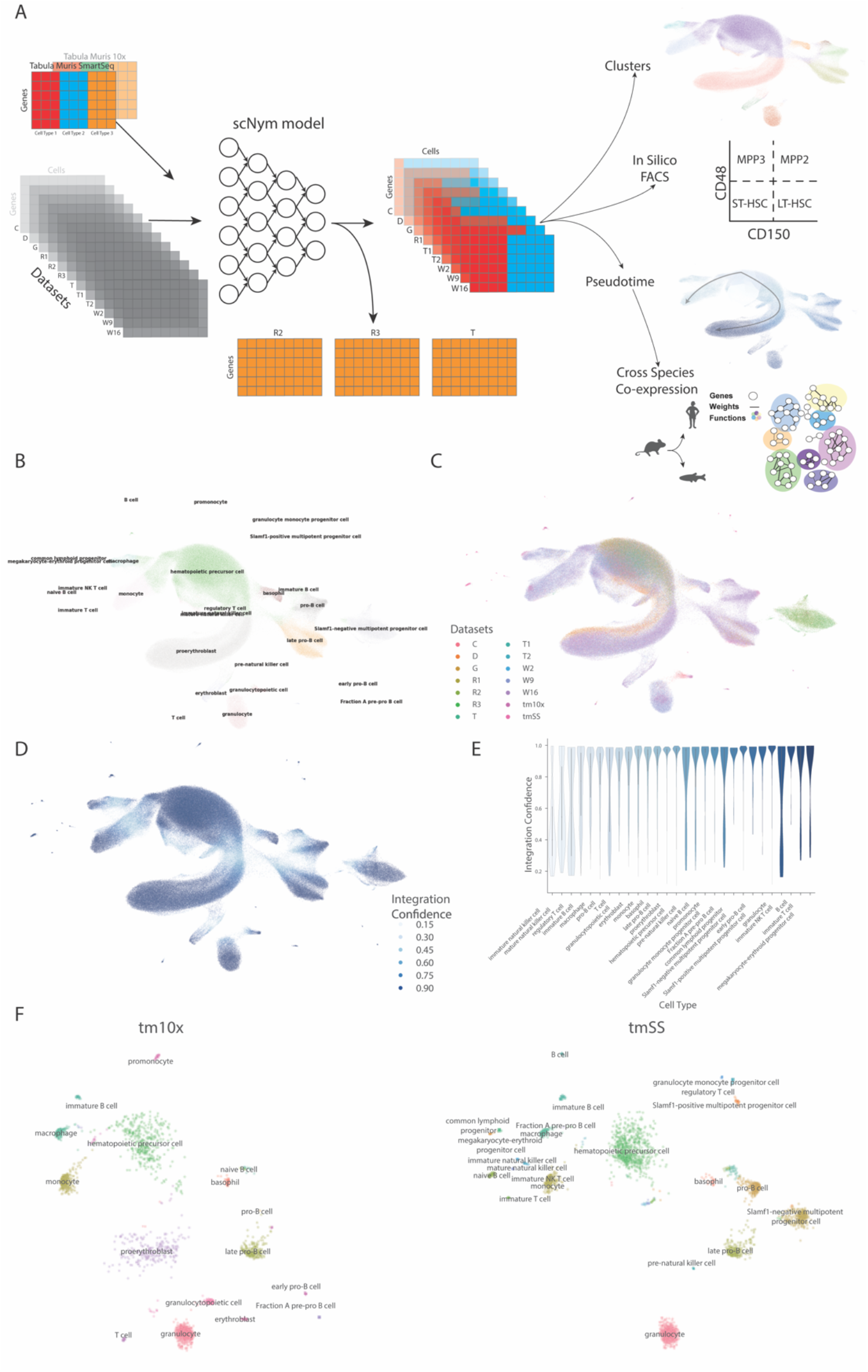
Integrations and filtering of unlabeled datasets using the Tabula Muris and scNym. **A)** Two tabula muris bone marrow datasets are used as references with scNym to label 12 datasets. 3 datasets are excluded from further analysis due to poor alignment with the remaining 9 datasets. The 9 remaining datasets are evaluated using a cluster, *in silico* FACS, and pseudotime analysis. The results of the psuedotime analysis are evaluated across many species. **B-C)** UMAP projection of the integrated datasets colored by **B)** cell type annotated by scNym and **C)** dataset. **D)** The confidence score for each cell type label in the UMAP projection. **E)** Confidence scores by cell type show most cells within a cell type are confidently labeled. **F)** UMAP projection of the reference tabula muris datasets show disconnected clusters.

It is important to assess cell type label accuracy when transferring cell type labels from reference data using the confidence scores computed by scNym. In the reduced space, the high confidence areas are cells towards the center of clusters, while lower confidence cells are in the areas between cluster centers (Figure 1D-E). Displaying just the training Tabula Muris data in the latent space makes it clear that the high confidence scores are in the regions of the latent space occupied by Tabula Muris cells, and regions of low confidence are in between the reference cell types (Figure 1F). The low confidence between clusters reflects the degree to which the model is extrapolating outside the training data space. Most of the Tabula Muris clusters are islands in the latent space with no adjoining neighbors. We suspect this is because the Tabula Muris cells were sorted based on cell surface markers to enrich for specific cell types before sequencing ^23^. This selects for more transcriptionally homogenous populations, useful for annotation but less so for understanding lineage relationships and variability. On the other hand, the datasets labeled by scNym were only sorted to broadly include hematopoietic stem and progenitors and some lineage-committed cells. After the integration, we removed the datasets R2, R3, and T because they did not map to the other 9 datasets’ cell types (Supplementary Figure 2A). We exclude the tabula muris datasets, R2, R3, and T to focus on datasets that sampled similar portions of the hematopoietic lineage for the remaining analysis.

### Robust Clustering

The remaining 9 datasets all broadly cover the same area of the latent space (Figure 2A-B, Supplementary Figure 2B). Most identified cell types are in all 9 datasets; 6 of the 13 clusters are shared across all datasets, and the remaining clusters are in at least 5 of the 9 datasets (Figure 2C). Every cell type contains at least one marker with an AUROC > .8 and a large fold change (Figure 2D, Supplementary Table 1). The top markers are very specific to the clusters they identify (Figure 2E). Klf1 and Ermap, two genes identified as markers for proerythroblast, are commonly known as erythroid markers ^34,35^. In our dataset selection process, we focused on studies that sorted cells based on the commonly used LSK markers: Lin-, Sca1+ (some Sca1-), and cKit+. The expression of cKit distinguishes proerythroblasts from more differentiated cell types in the erythroid lineage ^36^. Between the selection method and the marker genes, we are confident in the identification of the proerythroblast lineage, especially over more differentiated cell types within the lineage.

**Figure 2:**
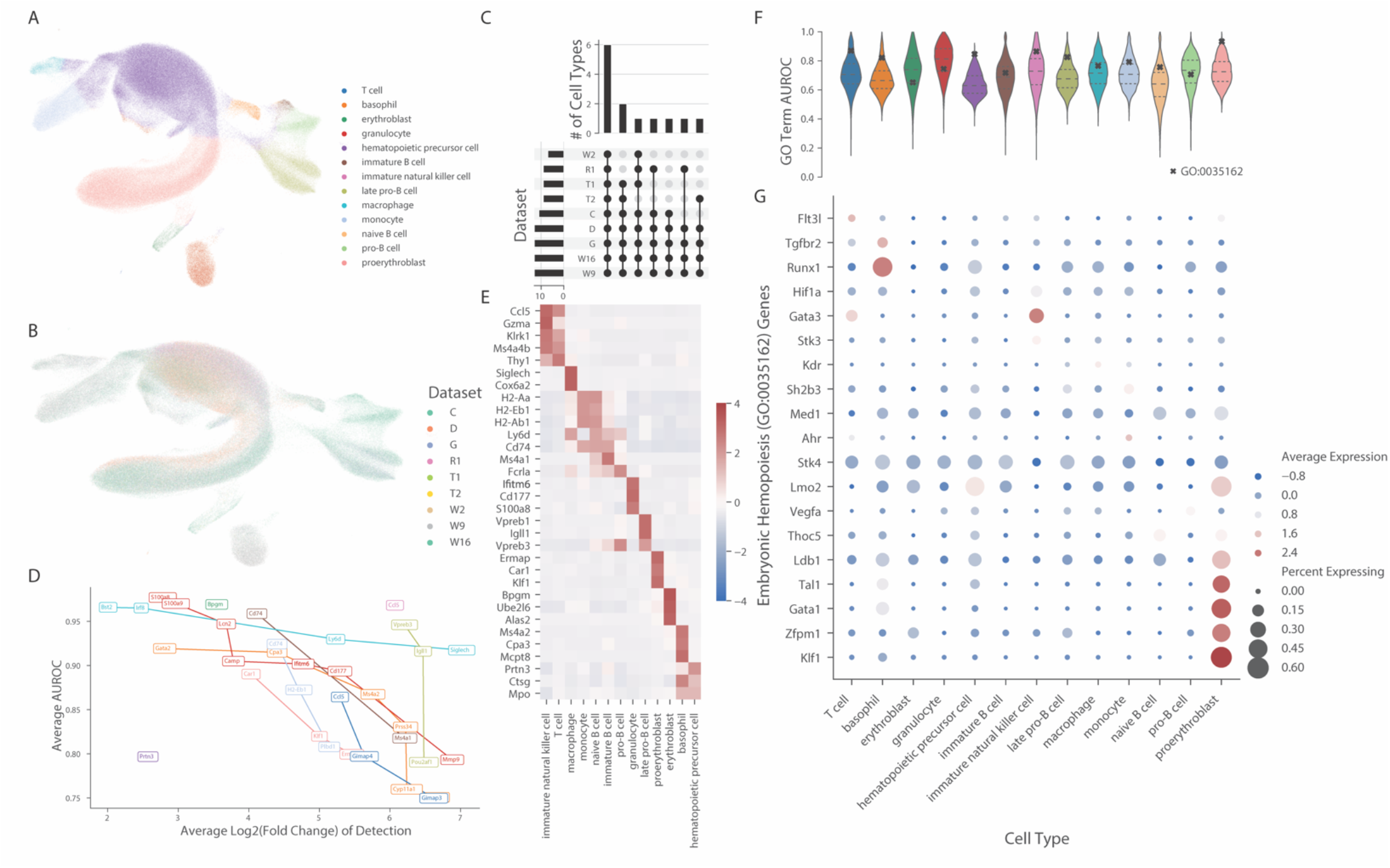
Identification of robust markers and functional programs that define cell types. **A-B)** UMAP projection of cells retained for downstream analyses colored by **A)** cell type label and **B)** dataset. **C)** Upset plot depicts that most cell types are shared across datasets. **D)** Each cell type has high-performing markers, as calculated using MetaMarkers. The top markers are plotted by their significance (Average AUROC) and effect size (Average Log Fold Change of Detection). They are colored by the same scheme as in A). **E)** The top 3 markers for each cluster show high expression specificity in a heatmap of expression. Z scores were calculated within datasets and then aggregated across datasets to account for technical variation between datasets. **F)** Violin plot of AUROCS for the MetaNeighbor results run on the Gene Ontology identifies many highly robust functional annotations that identify cell types. The AUROCs for the term Embryonic Hematopoiesis (GO:0035162) are marked for each cell type. **F)** Expression levels and % of cells expressed for the genes in the term Embryonic Hematopoiesis (GO:0035162) identify genes associated with the cell types that have the highest AUROCs from MetaNeighbor. The expression values are computed by Zscoring within datasets and then aggregating the values across datasets.

We evaluate the function of each cell type by using MetaNeighbor to identify replicable functional programs associated with each cell type, as labeled by the Gene Ontology. MetaNeighbor characterizes gene sets by their ability to “barcode” particular cell types via their expression profile. Each cell type has at least 75 GO terms with an AUROC > .9 (Figure 2F, Supplementary Table 2), meaning that the set of genes within that GO term is highly characteristic for a cell type and replicable in its expression profile. The term Embryonic Hemopoiesis (GO:00035162) has an average AUROC of .79, with moderate variation between the different cell types (Figure 2F). We visualize that variation between the cell types with a dotplot to show the expression of the genes within the term in each cell type (Figure 2G). The high performance of the term on basophils is largely driven by Runx1 expression (AUROC=.82). This is consistent with previous studies that show knockout of Runx1 reduces basophil’s found in bone marrow by 90% ^37^. Proerythroblasts are the highest performing cluster on the term (AUROC = .92). Gata1, and two genes associated with Gata1 expression, Klf1, and Zfpm1, are enriched in the proerythroblasts. The co-expression of Lmo2 and Ldb1 in proerythroblasts is consistent with results that show their role as maintainers of erythroid progenitor states and preventing further differentiation into the erythroid lineage ^38^. The marker genes and genes identified from GO enrichment show that we are predominately sampling proerythroblasts from the erythroid lineage.

### In Silico Sorting Identifies Latent Stem Cell States

Sorting cells based on cell surface marker protein expression is the established way of defining hematopoietic stem and progenitor cell types. We use the same marker genes, Slamf1 (CD150), Slamf2 (CD48), and Flt3 to sort the hematopoietic precursor cell cluster into Long term HSCs (LT-HSC), Short-term HSC (ST-HSC), and Multipotent Progenitors (MPP2-4) based on published guides ^12,39^. Interestingly, they do not appear to spatially organize in UMAP space, even when each dataset is individually projected onto a latent space (Figure 3A). Using MetaNeighbor to evaluate the replicability of the cell states, there is moderate replicability, especially with the MPP4 and LT-HSCs (Figure 3B). MetaNeigbhor does not identify a strong distinction between the MPP2 and MPP3 labeled cells, but they are distinct from the remaining cell states.

**Figure 3:**
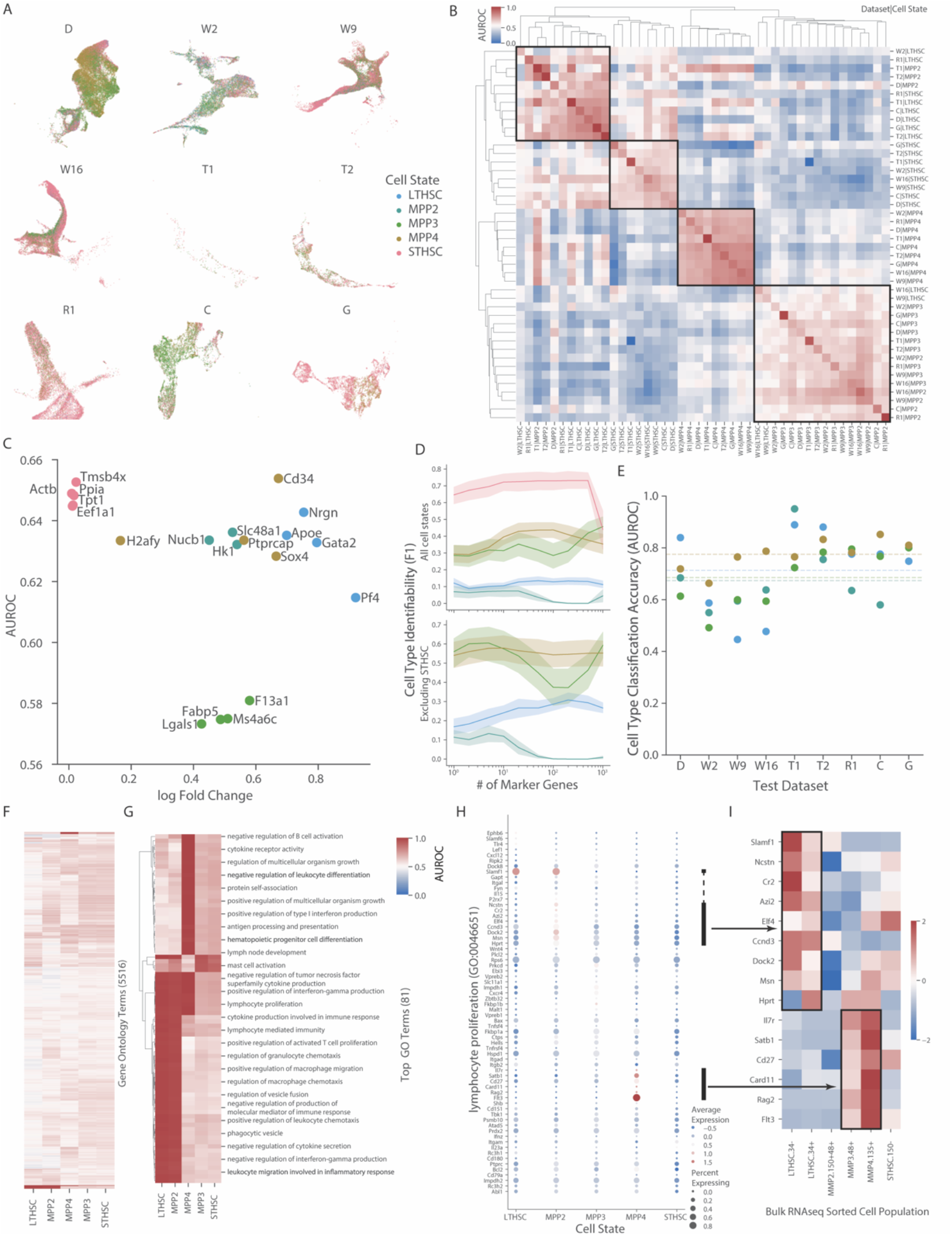
*in silico* FACS sorting identifies robust latent signatures of hematopoietic precursor cell states. **A)** Cells from the hematopoietic precursor cell labeled as either LT-HSC, ST-HSC, and MPP2-4 do not cluster in UMAP space when projected by dataset. **B)** MetaNeigbhor unsupervised analysis shows consistency of MPP4s across datasets and moderate replicability of the other cell states. **C)** MetaMarkers defined cell type markers show limited significance (AUROC) and weak effect sizes (log Fold Change). **D)** Evaluating the identifiability (F1 score) of cell states using 1-1000 markers in all cell states (top) and excluding the ST-HSC cell type (bottom). Computed using leave one out cross-validation. The shaded region represents 1 standard deviation. **E)** Classification performance (AUROC) using the top 10 marker genes. The model is trained on 9 datasets and the performance is shown for the 9th held out dataset. The dashed lines are the average across all folds. **F)** Metaneighbor evaluation of cell states using the whole Gene Ontology **G)** Subset of Metaneighbor results for terms with AUROC >.9 in at least 1 cell state **F)** Expression of the term lymphocyte proliferation (GO:0046651) in each of the cell states. The Z-scores are computed within datasets and then aggregated across datasets. **G)** Bulk expression from ImmGen data for genes with notable expression in the single cell data matches single cell expression.

The top marker genes show modest cell type predictability (AUROC) and weak signal-to-noise ratios (log Fold Change) (Figure 3C). The ST-HSCs have a near-even signal-to-noise ratio despite the highest predictability for the top markers. We test the identifiability of each cell state using the top 1-1000 markers to see if that does better than individual markers (Figure 3D). The ST-HSCs have modest identifiability, while the other cell states have extremely low identifiability (Figure 3D). ST-HSCs are the cell type defined by no expression of Slamf1, Slamf2 and, Flt3, given the sparsity of scRNAseq data and the low signal to noise ratio for ST-HSC marker genes, it could be possible that the cell type is a mixture of actual ST-HSCs and the other cell states incorrectly labeled. When removing the ST-HSCs, the identifiability (F1) increases to moderate levels for the MPP3 and MPP4 cell states using as few as 10 markers. LT-HSC identifiability is extremely low with 1 gene but steadily increases with the number of markers. To look at the variation across datasets we learn the top 10 markers for each cell state in 8 datasets and measure how well they classify the held out (test) dataset. The average AUROC across all the tests is .71, but with considerable variability between the different datasets and cell states (Figure 3E). Classifying these cell states across datasets provides modest performance. The MetaNeighor and marker classification analysis identify replicable axes of variation, even if not the primary ones that would be visible in UMAP space.

We evaluate the replicability of functional connectivity of gene sets within the cell states using MetaNeighbor. Most of the 5516 tested GO terms have consistently low AUROCs across all cell states (Figure 3F, Supplementary Table 3). However, 81 terms have an AUROC >.9 in at least 1 cell state (Figure 3G). Within top enriched terms, we see that many match the known differentiation bias of MPPs. The GO term “Lymphocyte Proliferation” (GO:0046651) has an AUROC of .98 in the MPP4 cluster. MPP4s are also referred to as the Lymphoid Multipotent Progenitor (LMPP) and have a significant bias towards differentiating into the lymphoid lineage 2. The expression patterns for the genes in the term are displayed in a dotplot (Figure 3H). The most variably expressed genes in the term show expression patterns consistent with bulk sorted cell populations from the Immgen Consortium (Figure 3I, ^40^). Rag2 and Il7r are standard markers for B and T cell development and Satb1 promotes lymphocyte differentiation ^41^. The enrichment of the lymphoid proliferation term and lymphoid-associated genes could indicate that the cells in the MPP4 cell state are lymphoid primed. While not the primary axis of variability, these cell states constitute a replicable axis of variation within the hematopoietic precursor cell cluster associated with lineage potential.

### Robust Signatures of Hematopoietic Differentiation

Modeling the cells as an ordered continuum, instead of clusters, depicts the differentiation process within the data and can identify gene regulation dynamics specific to lineage determination. We model this by computing pseudotime in individual datasets to avoid learning trajectories that are artifacts of the integration process/batch effects (Figure 4A). The pseudotime computed on the integrated space is markedly different for each dataset (Supplementary Figure 3). In addition to producing an ordering of the cells, the algorithm assigns all of the cells to nodes along a tree that estimates the differentiation branching within the data. We associated each end segment of the trees to either root, erythroid, monocyte based on gene expression and label all segments in the middle as intermediate. While the clustering includes lymphocyte cells, the individual dataset projections do not connect the lymphocyte cells to the root in the latent space and we can not compute a confident trajectory through the non-linear gaps in the latent space (Supplementary Figure 4). Evaluating the replicability of the segments using MetaNeigbhor shows that the root, erythroid and monocyte segments are replicable across the datasets, while the intermediate segments are not replicable across the datasets (Figure 4B, Supplementary Figure 5). The inconsistency of the intermediates could be a result of the transient nature of intermediate cell types or more technical issues with scRNAseq ^42,43^.

**Figure 4:**
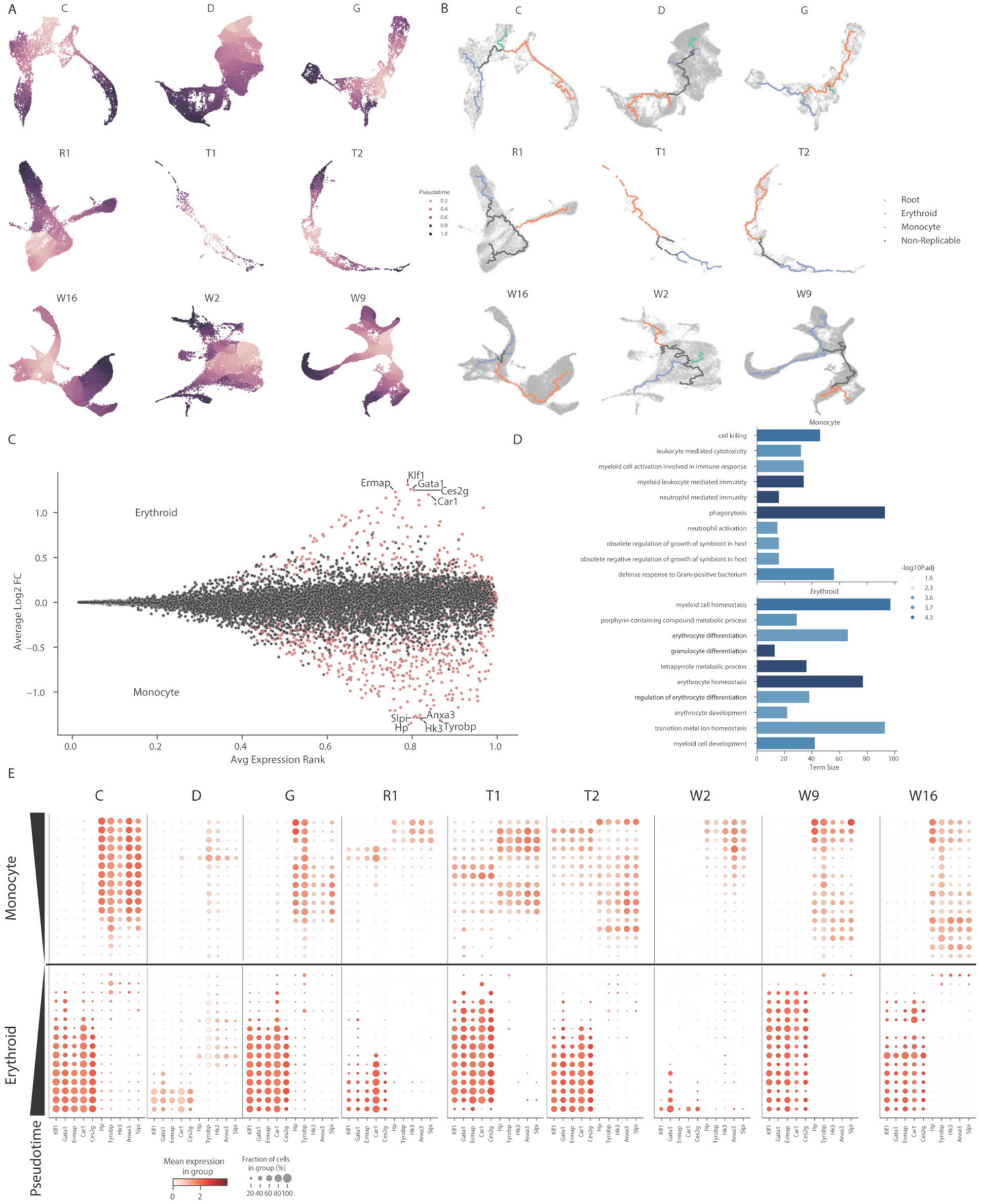
Pseudotime analysis in individual datasets identifies lineage-associated modules and sampling differences between datasets. **A)** Individual datasets are projected into 2-dimensional space using UMAP and then Monocle3 learns a pseudotime ordering of the cells. **B)** Branches of the pseudotime trajectories are assigned to either Root, Erythroid, Monocyte, or Non-replicable based on MetaNeighbor results (Supplementary Figure 5). **C)** Meta-analytic MAplot of marker genes for Erythroid and Monocyte lineages. **D)** Top 10 terms from Gene Ontology enrichment using fisher’s exact test for the top 50 makers for each lineage. **E)** Expression of top 5 markers from each lineage across datasets ordered by pseudotime shows monotonic patterns but different expression profile dynamics between datasets.

Using a broader approach, we fit models for every gene to each dataset and use meta-analytic statistics to identify consistent gene expression signatures associated with the erythroid and monocyte lineages (Figure 4C, Supplementary Table 4). The top 3-5 genes for lineages are very similar to the cluster level analysis for erythroid, but not for monocyte. However, looking at the top 50 genes for each dataset shows that only 19 for erythroid and 4 for monocyte genes are shared between the cluster and pseudotime analysis (Supplementary Figure 6). GO enrichment of the top 50 genes for each lineage identifies 11 for erythroid and 54 for monocyte significantly associated terms (p<.05, Figure 4D, Supplementary Table 5). Visualization of the top 5 genes for each lineage ordered by pseudotime shows a consistent monotonic expression trend across the datasets (Figure 4E). Despite the consistent monotonicity, each dataset has a unique inflection point where the gene expression substantially increases. The differences in timing across the datasets explain some of the replicability limitations of comparing the intermediate cells across datasets.

### Cross-Species Co-Expression of Lineage Signatures

Co-expression networks reflect the functional landscape of gene expression ^44^. Reference, bulk-RNAseq derived, co-expression networks are used to evaluate the cross-species relevance of the lineage-associated gene lists ^45^. We measure the connectivity (AUROC) of the erythroid and monocyte gene lists using these co-expression networks (Figure 5A). strong connectivity, or high AUROC, of a gene set indicates shared function. As expected, the highest co-expression for both gene lists is in the mouse network; the training species for the gene lists (monocyte AUROC=.92, erythroid AUROC=.90). Using 1-to-1 orthologs we evaluate the co-expression of the gene lists in 21 species. The monocyte gene list is more co-expressed in most species than the erythroid gene set. At the extreme is zebrafish, with near-random co-expression for erythroid (AUROC=.42) and strong co-expression for monocyte genes (AUROC=.81). Strikingly, both gene sets perform well in the human co-expression network, indicative of strong mouse-human conservation, an encouraging sign for translational research purposes (monocyte AUROC=.88, erythroid AUROC=.82).

**Figure 5:**
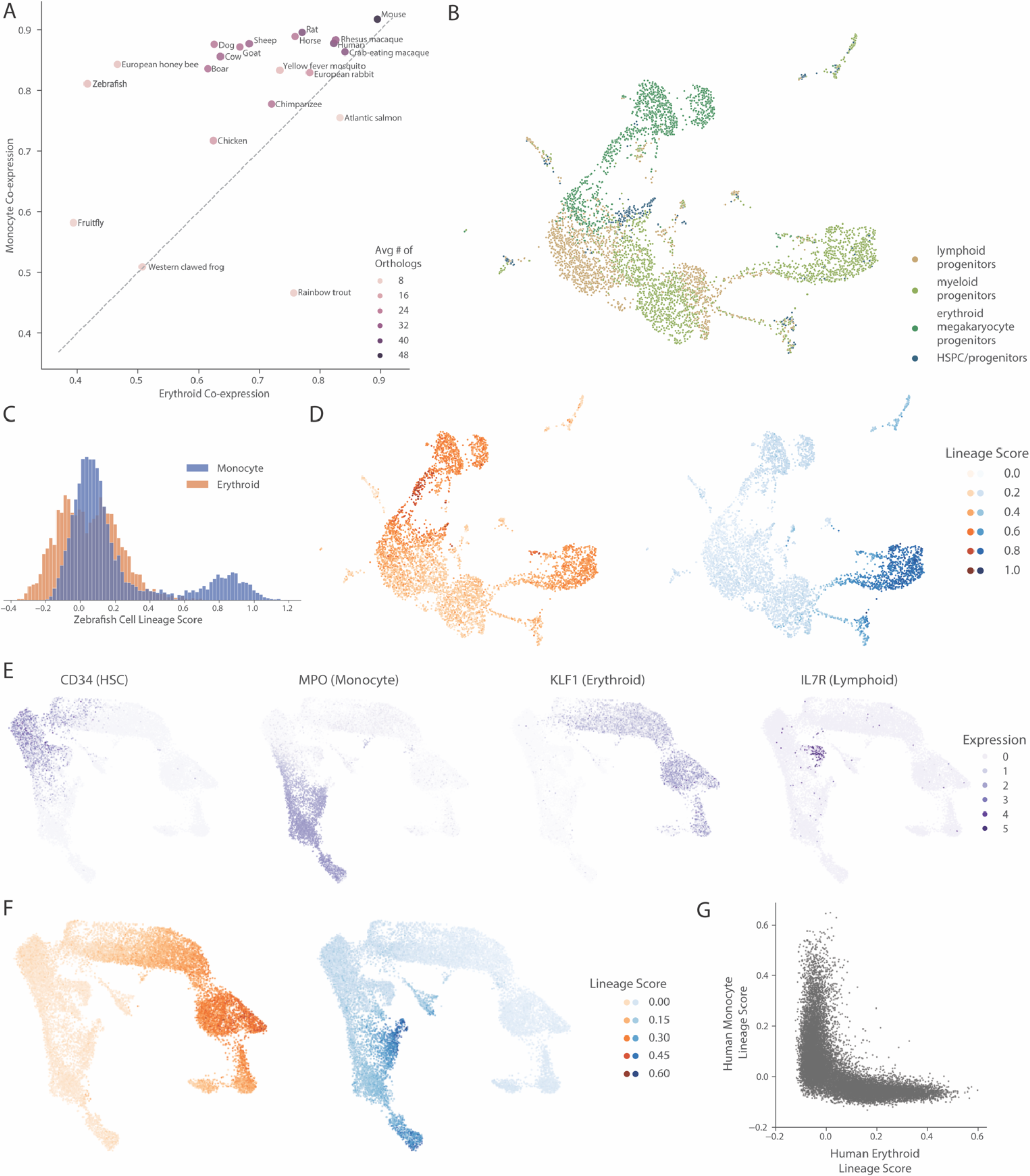
Cross-species co-expression analysis predicts functional conservation of lineage-associated gene sets. **A)** Co-expression of 1-to-1 orthologs across 21 species for both the erythroid and monocyte associated gene lists shows bias towards conservation of the monocyte lineage. **B)** UMAP of Xia et al 2021 zebrafish hematopoietic dataset colored by cell type label. **C)** Histogram of lineage scores for each cell in zebrafish dataset has enriched population of cells for only the monocyte gene list **D)** Lineage score for both monocyte and erythroid lineages plotted on UMAP of dataset shows only has specificity for monocyte lineage. **E)** Expression of known markers for major hematopoietic lineages in the human hematopoietic dataset Pellin et al 2019. **F)** Scores for erythroid and monocyte lineages in the human dataset specifically identify both erythroid and monocyte cell populations **G)** The lineage scores for each cell in the human dataset show the gene programs are orthogonal to each other.

In addition to evaluating conservation using co-expression networks, we look at the expression of the gene sets in a zebrafish hematopoietic dataset (Figure 5B, ^46^). The monocyte scores are bimodal, with the highest scoring cells matching the cells labeled as myeloid progenitors in the original study (Figure 5C-D). They mostly have very low scores for the erythroid gene set, and many of the highly scoring cells are myeloid progenitor-labeled cells. We next assessed the human bone marrow dataset from Pellin et al 2019 to evaluate the expression of the gene sets in human data ^47^. We rely on top markers from the publication to identify the HSCs, Monocyte, Erythroid, and Lymphocyte cell populations because discrete labels were unobtainable (Figure 5E). The scores form increasing gradients from the HSCs to their respective lineage (Figure 5F). The two lineage scores are orthogonal to each other, showing they serve as a good marker for the lineages (Figure 5F). The orthogonal signatures show a binary fate decision between the erythroid and monocyte lineages; if one lineage score is upregulated, the other one remains inactivated. Between the co-expression and human scRNAseq results, it is clear that the functional relationship of the genes in the lineage-associated gene lists is conserved between mice and humans.

## Discussion

Our results provide a robust evaluation of hematopoietic cell populations in mouse bone marrow. After identifying 9 hematopoietic datasets that broadly share cell types, we identified cellular populations using 3 different methods: clustering, in silico FACS sorting, and trajectory inference. These populations were characterized using both markers and functional annotations. Furthermore, we demonstrated the conservation of lineage-associated genes using co-expression analysis across 21 species. Finally, we made the data and identified signatures accessible on our shiny webserver to compare with future experiments.

Meta-analysis serves to find robust signatures across datasets with significant technical variation ^15^, thereby determining what markers and properties are likely to generalize to new data. This meta-analytic atlas resolves technical limitations with individual batches to better represent the continuous nature of the system and provide strongly replicable signatures. The datasets sample cells unbiasedly from hematopoietic progenitors, recapitulating a developing system unlike the discrete, FACS sorting-based sampling in the Tabula Muris ^23^. The most popular present resource for hematopoietic transcriptomic signatures is built from a single bulk RNAseq dataset, but has still been invaluable for basic research and studying SARS-Cov2, tuberculosis, and leukemia ^40,48–53^. By extending the availability of reference data to single cell and comparing across datasets, we enhance both the depth and breadth of transcriptomic signatures available to researchers.

The generalizability of our results will make it a valuable resource for translational research. An accurate reference of healthy hematopoietic stem cells is critical for identifying reliable therapeutic targets. While learning functional signatures of disease from clinical samples is often preferable, they can be difficult to acquire, and an alternative is to learn signatures associated with diseases from mouse models ^54,55^. In order to identify disease signatures, correctly identifying cell types in healthy conditions is critical for evaluating changes in expression or abundance. Disease-associated signatures identified in single-cell data could then be evaluated as arising from changes in expression within cell types, or changes in cell type proportions ^56,57^. Importantly, our cross-species analysis shows that we can evaluate the conservation of signatures identified in mice to human data, demonstrating the atlas’ utility for pre-clinical therapeutic research.

Here, we focus on the integration of one data modality, scRNAseq, but we expect additional modalities to be incorporated as data continues to be generated and robust meta-analysis can be conducted. In general, expression data serves as a foundation for the integration of other data modalities, providing robust signatures which can then be annotated by the data used in other modalities ^18^. A cross-dataset, multi-modal atlas will resolve limitations and produce a more detailed picture of the gene regulatory networks driving hematopoiesis. Integrating CITE-seq data, which measures cell surface protein expression and RNA with this atlas will resolve the progenitor states better than *in silico* FACS sorting ^58^. Single cell ATACseq data from mouse bone marrow will identify transcription factors and cis-regulatory elements important to lineage commitment ^59^. CRISPR screens will test lineage-specific gene dependencies ^60,61^. Cell non-autonomous signaling influences lineage commitment, either from the non-hematopoietic cells in the bone marrow or cell-cell communication between hematopoietic populations ^62–64^. Evaluating such cell-cell interactions will identify external signals that dictate lineage commitment. More data covering gaps in continuity, particularly the lymphoid lineage, will generate a more complete atlas— of great utility for studying lymphoid malignancies. Integrating other modalities with our robust scRNAseq atlas will resolve gaps in the atlas and produce a high-resolution picture of hematopoietic development.

This atlas serves as a reference for future hematopoiesis experiments that transition from FACS, the current gold standard, to RNA expression as the phenotypic measurement. In our results, we demonstrate multiple targeted analyses, made possible by a meta-analytic atlas and web server. Our analysis provides a detailed and robust evaluation of hematopoietic lineage development in mouse bone marrow. Our webserver makes it easy to evaluate the expression of any gene or known function identified in future experiments (https://gillisweb.cshl.edu/HSC_atlas/).

## Acknowledgments

B.D.H. was supported by the CSHL Crick Cray Fellowship. J.G. and J.L. were supported by NIH grants R01MH113005 and R01LM012736. This work was performed with assistance from the US National Institutes of Health Grant S10OD028632-01. The authors would like to thank Diogo Maia-Silva for their useful input and discussion about the experiments and manuscript.

## Author Contributions

B.D.H. and J.G. conceived the experiments and prepared the manuscript. B.D.H. conducted all of the experiments. J.L. developed and maintains the webserver.

