## Supplemental Figures for "A Meta-Analytic Single-Cell Atlas of Mouse Bone Marrow Hematopoietic Development"

### Supplementary Figures

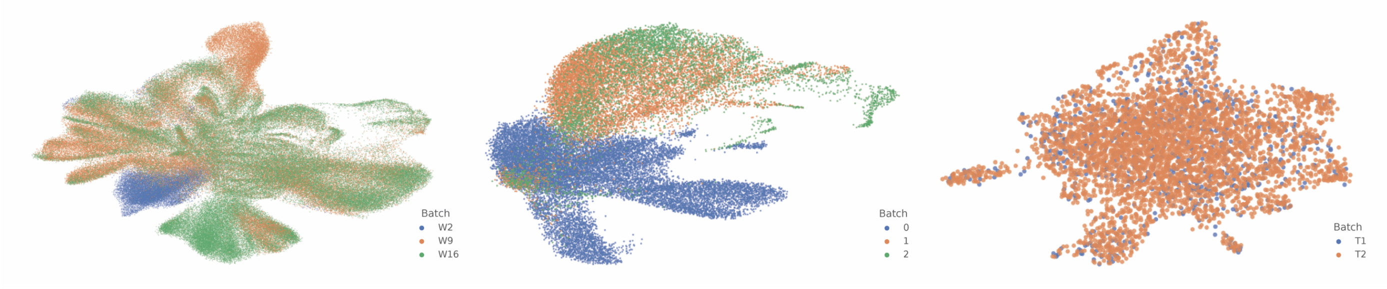


**Supplementary Figure 1: Evaluating the visibility of batch effects from 3 publications.** Projecting the datasets Weinreb et al 2020 (left), Rodriguez-Fraticelli et al 2020 (middle), and Tikhonova et al 2019 (left) using UMAP and colored by the batch label from the metadata shows strong batch effects in the left and middle datasets.

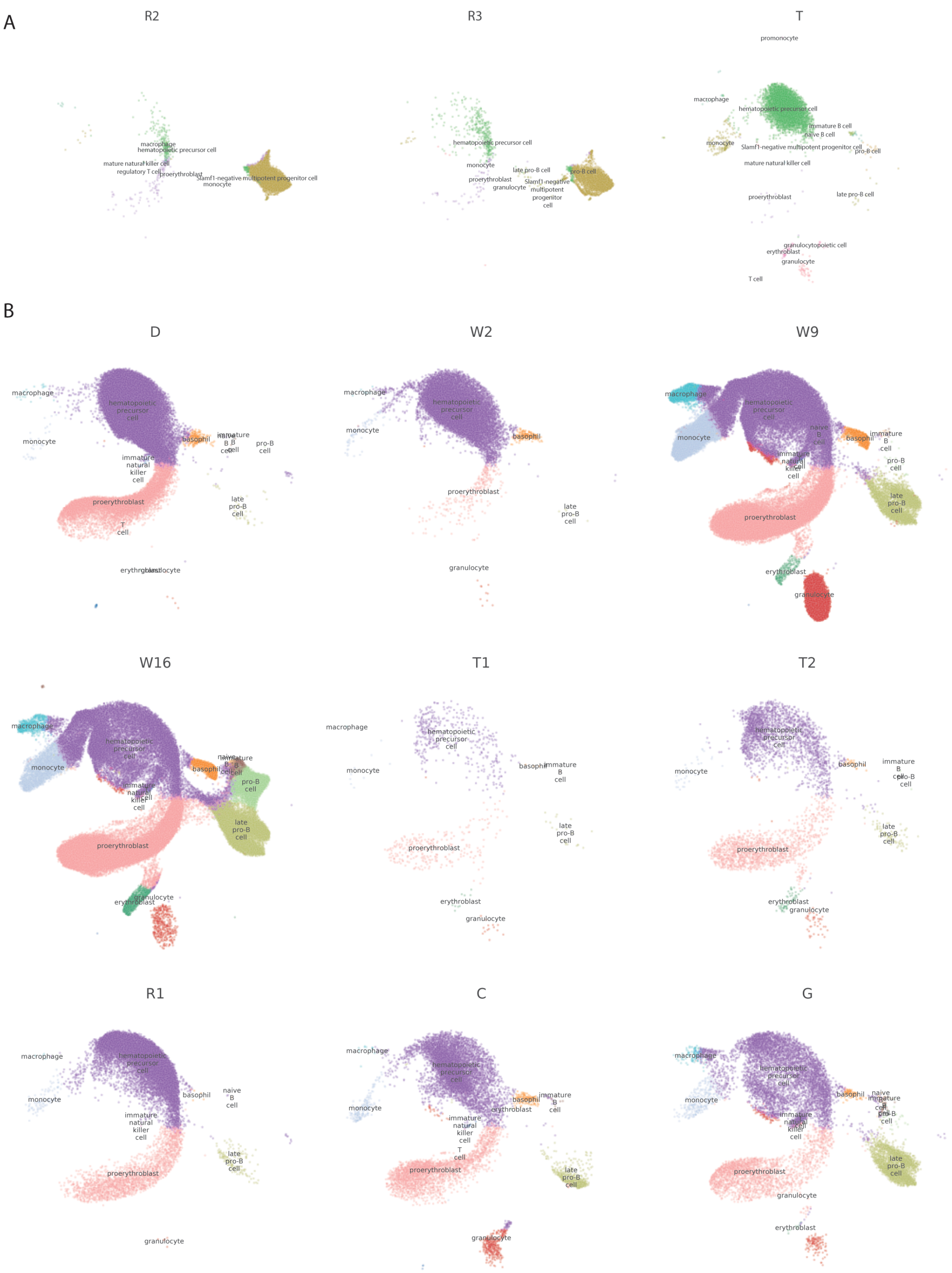


**Supplementary Figure 2: UMAP projection of the individual integrated datasets. A)** Projections in the integrated latent space of the 3 datasets excluded from downstream analysis colored by cell type **B)** Projection in the integrated latent space of the 9 datasets that share the same cell types and cover the same region of the integrated latent space

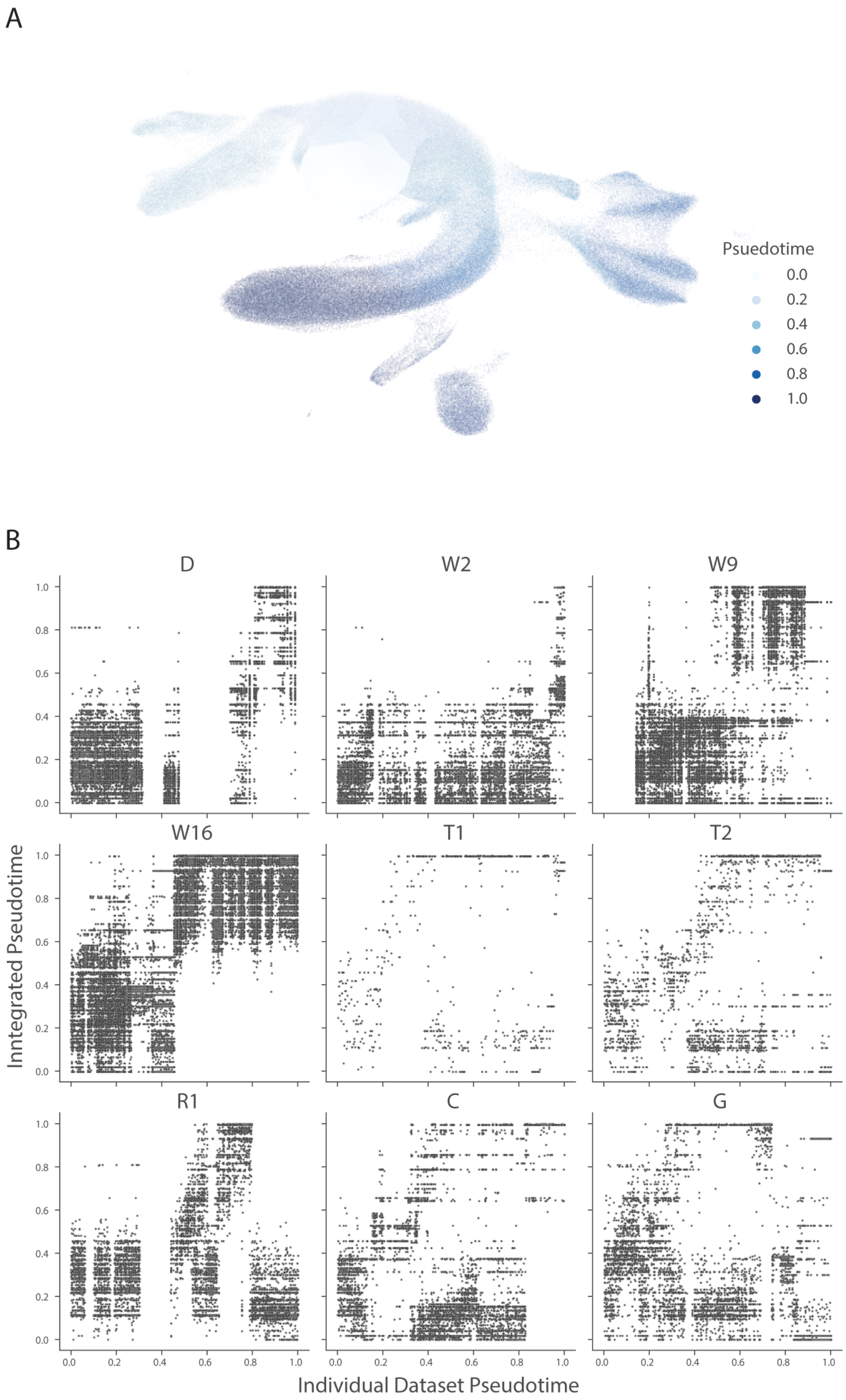


**Supplementary Figure 3: Pseudotime ordering of integrated space produces a different ordering of the cells than pseudotime within individual datasets. A)** Ordering of cells computed by Monocle3 for the integrated latent space, only including cells that were used in the individual dataset pseudotime analysis. **B)** Pseudotime ordering of individual datasets compared is different from the integrated ordering

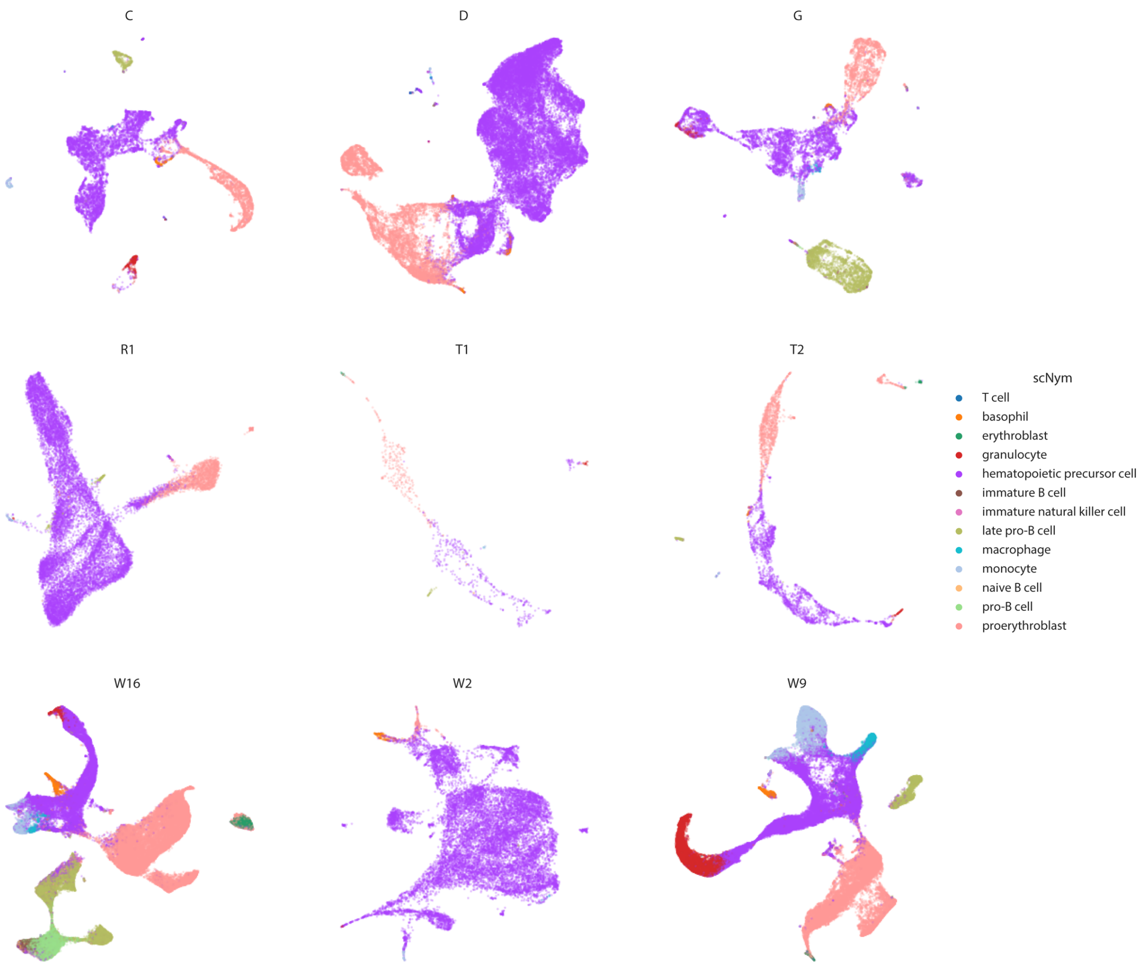


**Supplementary Figure 4: Projection of individual datasets shows disconnected lymphoid lineage.** Projecting individual datasets using UMAP shows the lymphoid lineage (B+T cells) are disconnected from the stem cell and other clusters. Computing a trajectory between disconnected clusters is ill-advised.

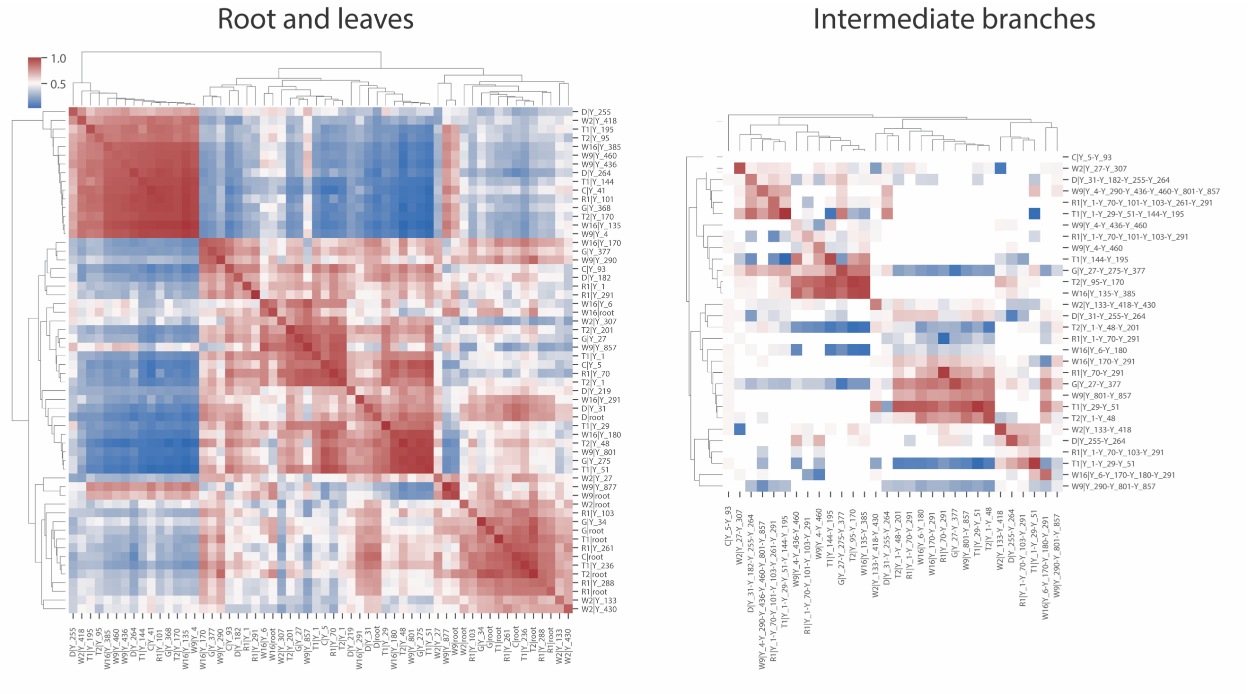


**Supplementary Figure 5: Unsupervised MetaNeighbor identifies replicable root and erythroid branches and non-replicable intermediate branches**

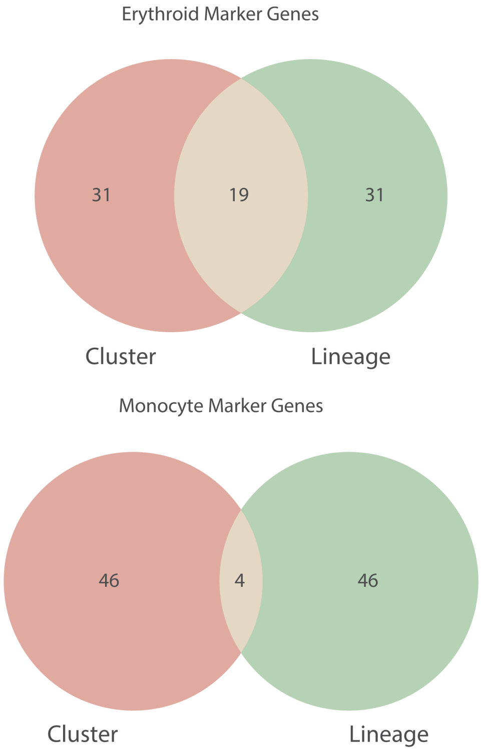


**Supplementary Figure 6: Modest overlap between the top 50 markers from cluster-level analysis and pseudotime analysis.**
